## Supplemental Materials for "Statistical crystallography reveals an allosteric network in SARS-CoV-2 M^pro^"

Creon *et al.*

**This PDF file includes:**

Figs. S1 to S11
Tables S1 to S5
Supplemental References

**Other Supplementary Materials for this manuscript include the following:**

Data S1 to S5

S1. SEC raw data
S2. SEC fit data
S3. ITC and nDSF raw data
S4. Raw reaction velocities
S5. Fit Michaelis-Menten parameters

**Supplemental Table 1. Crystallographic statistics for the N214A, Q256A, and S284A mutants.** Statistics for the highest-resolution spherical shell are shown in parentheses.

| $M^{pro}$ | N214A | Q256A | S284A |
|---|---|---|---|
| **PDB ID** | 9GI6 | 9GHN | 9GHO |
| **Data collection** | | | |
| Source | PETRA-III | PETRA-III | PETRA-III |
| Temperature | 100 K | 100 K | 100 K |
| Space group | $P2_12_12_1$ | $P2_12_12_1$ | C2 |
| *a*, *b*, *c* (Å) | 67.81 100.57 102.18 | 78.54 88.44 100.77 | 113.20 53.27 44.84 |
| α, β, γ (°) | 90 90 90 | 90 90 90 | 90 102.4 90 |
| Resolution Cut | anisotropic | isotropic | isotropic |
| Resolution (Å) † | 55.28 – 2.51<br>55.28 – 2.58<br>55.28 – 1.83<br>(2.01 – 1.83) | 66.47 – 1.40<br>(1.45 - 1.40) | 55.28 – 1.86<br>(1.93 – 1.86) |
| Wilson B (Å$^2$) † | 45.29 77.34 58.51 | 19.09 | 36.44 |
| $R_{merge}$ | 0.080 (0.400) | 0.049 (1.505) | 0.124 (1.835) |
| $R_{meas}$ | 0.086 (0.438) | 0.053 (1.649) | 0.144 (2.134) |
| $R_{pim}$ | 0.031 (0.221) | 0.020 (0.6602) | 0.071 (1.071) |
| $I / \sigma I$ | 12.1 (1.5) | 19.05 (1.05) | 4.68 (0.30) |
| $CC_{1/2}$ | 0.997 (0.943) | 0.999 (0.584) | 0.995 (0.179) |
| Total Reflections | 320 497 (15 980) | 992 672 (78754) | 83 255 (8213) |
| Unique Reflections | 42 912 (2156) | 137736 (13221) | 21 743 (2154) |
| Multiplicity | 7.5 (7.4) | 7.2 (6.0) | 3.8 (3.8) |
| Completeness (%) | 94.8 (64.5) | 99.42 (95.29) | 91.2 (65.1) |
| **Refinement** | | | |
| $R_{work}$ | 0.174 (0.217) | 0.193 (0.3761) | 0.196 (0.385) |
| $R_{free}$ | 0.207 (0.304) | 0.218 (0.4038) | 0.241 (0.454) |
| No. atoms | 5229 | 5537 | 2458 |
| Protein | 4755 | 4883 | 2366 |
| Ligand/ion | 164 ‡ | 16 | 9 |
| Water | 310 | 638 | 83 |
| *B*-factors | 36.26 | 25.53 | 41.77 |
| Protein (Å$^2$) | 35.50 | 24.50 | 41.77 |
| Ligand/ion (Å$^2$) | 52.28 | 43.74 | 51.01 |
| Water (Å$^2$) | 39.41 | 33.03 | 40.70 |
| r.m.s. deviations | | | |
| Bond lengths (Å) | 0.007 | 0.005 | 0.007 |
| Bond angles (°) | 0.84 | 0.83 | 1.03 |

† *staraniso* produces an elliptical diffraction limit (*1*); for the N214A dataset, we report the resolution cutoff and Wilson B factors along the three principal elliptical axes; statistics take this elliptical truncation into account
‡ 22 PEG chains (7 atoms/chain) were modeled into density observed in the N214A dataset. See *Materials and Methods* for crystallization conditions of each variant.

| dataset | cell volume (nm$^3$) | ensemble size | single $R_{work}$ | single $R_{free}$ | ensemble $R_{work}$ | ensemble $R_{free}$ |
|---|---|---|---|---|---|---|
| **A** | 256.28 | 67 | 0.1997 | 0.2378 | 0.1566 | 0.2126 |
| **B** | 260.66 | 50 | 0.1791 | 0.2058 | 0.1577 | 0.2075 |
| **C** | 264.53 | 29 | 0.1955 | 0.2342 | 0.1693 | 0.2250 |
| **D** | 270.65 | 50 | 0.1870 | 0.2173 | 0.1627 | 0.2174 |

**Supplemental Table 2. Ensemble refinement statistics**. Ensemble refinements were performed on datasets selected to span the observed range of cell volumes to increase the diversity of the data analyzed. Reported are the number of conformations that *phenix* automatically determines during the ensemble refinement procedure, as well as *R*-factors for the single-conformer starting model and multi-conformer ensemble after MD-based refinement (*2*, *3*). These ensembles were used to compute covariance matrices, shown in Supplemental Figure 4; the first column maps these datasets to the panels shown there.

| variant | [E] (µM) | $K_D$ (µM) | *ΔH* (kcal/mol) | *ΔG* (kcal/mol) | *-TΔS* (kcal/mol) |
|---|---|---|---|---|---|
| wild type | 20.7 +/- 1.2 | 2.4 +/- 0.8 | -3.9 +/- 0.3 | -7.8 | -3.9 |
| N214A | 17.5 +/- 0.6 | 2.0 +/- 0.4 | -4.4 +/- 0.2 | -7.9 | -3.5 |
| Q256A | 17.5 +/- 1.3 | 3.4 +/- 1.0 | -5.9 +/- 0.6 | -7.5 | -1.7 |
| S284A | 16.7 +/- 0.7 | 1.6 +/- 0.4 | -3.0 +/- 0.2 | -8.0 | -5.1 |

**Supplemental Table 3. Thermodynamic parameters of calpeptin binding to wild type, N214A, Q256A and S284A M$^{pro}$ derived from isothermal titration calorimetry (ITC).** Enzyme concentrations ([E]), dissociation constants ($K_D$), enthalpy changes (*ΔH*), Gibbs free energy changes (*ΔG*), and entropy contributions (*-TΔS*) for each variant:calpeptin interaction were estimated from the standard two-state model fit to ITC data (Supplemental Fig. 6). Fits performed with MicroCal PEAQ-ITC. The comparable $K_D$ values suggest similar ligand binding affinities across the different mutants. Errors are standard errors of the mean, as reported by MicroCal PEAQ-ITC v1.41.

| **variant** | **[E] (μM)** | ***n*** | **$K_D$ (μM)** | **fit method** |
|---|---|---|---|---|
| wild type | **2** | **3** | **0.5 +/- 0.2** | **averaging** |
| | 10 | 3 | 0.4 +/- 0.2 | averaging |
| | combined | 6 | 0.43 +/- 0.12 | least squares |
| N214A | **2** | **3** | **5.6 +/- 0.9** | **averaging** |
| | 10 | 3 | 6 +/- 3 | averaging |
| | 20 | 3 | 4 +/- 3 | averaging |
| | combined | 9 | 5.9 +/- 1.4 | least squares |
| Q256A | **2** | **3** | **0.80 +/- 0.11** | **averaging** |
| | combined | 3 | 0.80 +/- 0.14 | least squares |
| S284A | **2** | **3** | **0.56 +/- 0.11** | **averaging** |
| | combined | 3 | 0.57 +/- 0.13 | least squares |

**Supplemental Table 4. Dimerization constants determined by native mass spectrometry.** To investigate the possibility of systematic errors as a function of concentration, monomer/dimer $K_D$s were computed from native mass spectrometry data one concentration at a time via simple averaging, and in aggregate by fitting the standard two-state law of mass action with non-linear least squares. Within error, the determined affinities are consistent. As low concentration measurements are expected to suffer the least systematic error, affinities determined at 2 μM (bold) were reported in the main text and employed in subsequent analysis. All uncertainties are standard errors assuming a Gaussian error model.

| model | (ii) third order | | (iii) dimer pre-equilibrium | | |
|---|---|---|---|---|---|
| **variant** | $k_{cat}$ (min$^{-1}$) | $K_{eq}$ (μM$^2$) | $k_{cat}$ (min$^{-1}$) | $K_M$ (μM) | $K_D$ (μM) |
| wild type | 1.6 +/- 0.4 | 1000 +/- 700 | 0.88 +/- 0.2 | 107 +/- 3 | 0.30 +/- 0.02 |
| N214A | 0.11 +/- 0.06 | 10 000 +/- 13 000 | 0.100 +/- 0.005 | 220 +/- 10 | 4e-26 +/- 8e-6 |
| Q256A | 0.8 +/- 0.2 | 900 +/- 500 | 0.32 +/- 0.02 | 54 +/- 3 | 0.18 +/- 0.04 |
| S284A | 2.1 +/- 0.6 | 2000 +/- 1000 | 0.50 +/- 0.01 | 57 +/- 2 | 0.019 +/- 0.005 |

**Supplemental Table 5. Fit parameters for alternative kinetics models.** See *Materials and Methods* for model details and Supplementary Figure 8 for a visualization of the data and fits. All errors are 95% confidence intervals.

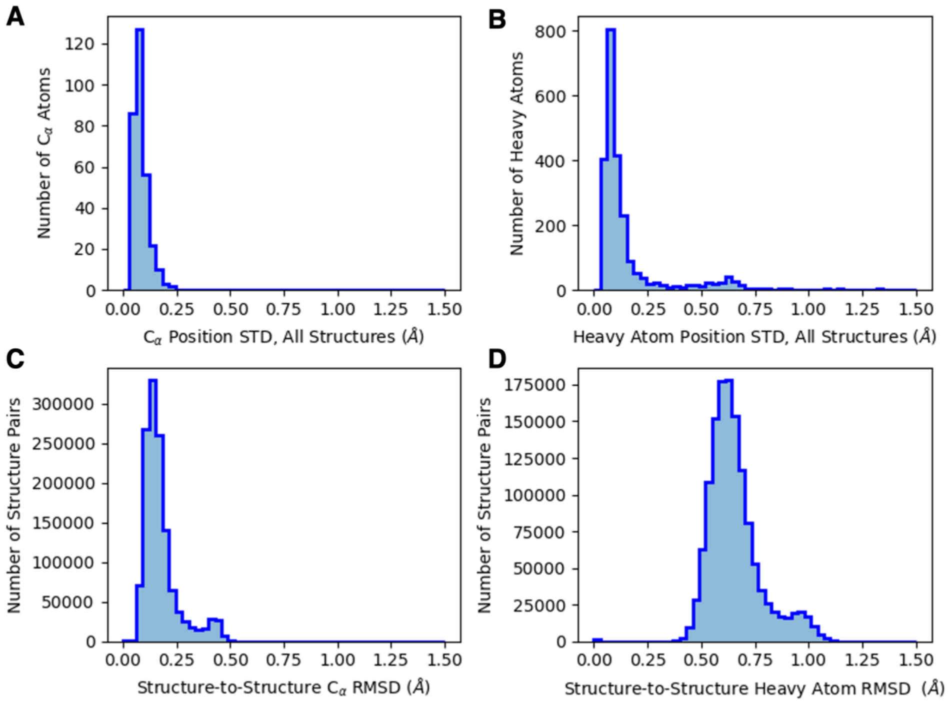


**Supplemental Figure 1. Summary of refinement results for the 1146 structure ensemble.** (**A**, **B**) The standard deviation of the refined atom positions provides a measure of the structural variability in the dataset produced by the automatic refinement procedure. Shown are (**A**) $C_\alpha$ atoms only and (**B**) all heavy atoms. Similarly, (**C**, **D**) the distribution of all pairwise structure RMSD values, again for (**C**) $C_\alpha$ atoms only and (**D**) all heavy atoms. The structural distribution is narrow, but non-trivial; clear bimodal behavior, for example, is apparent in this simple representation.

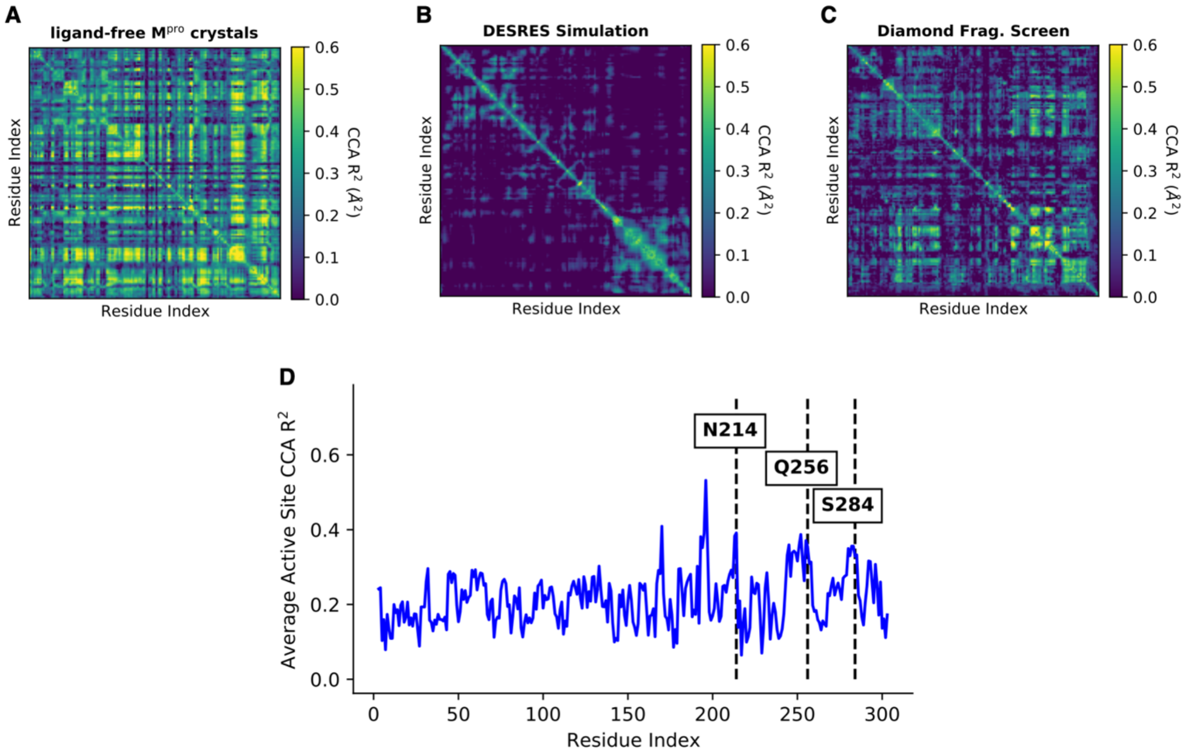


**Supplemental Figure 2. Canonical correlation analysis of the correlated motions in M$^{pro}$.** Given a pair of atoms (here C$_\alpha$), the canonical correlation reports the maximum correlation between any two linear functions of the Cartesian coefficients of each atom (x, y, z) (*4*). This differs from the covariance presented in the main text in two key ways. First, it accounts for situations where atomic correlations are non-isotropic, for example should the *x*-coordinate of atom *i* influence the *y*-coordinate of atom *j*. Second, it computes a correlation rather than a covariance, and therefore normalizes for the total magnitude of the atomic motion. Reported here are the coefficients of determination ($R^2$) for the atom-atom canonical correlation models of (**A**) our ensemble of 1146 crystal structures, (**B**) the DESRES MD simulation, and (**C**) the Diamond M$^{pro}$ structures, as presented in main text Figure 2. Further, shown is (**D**) the coefficients of determination averaged over active site residues, analogous to main text Figure 3C.

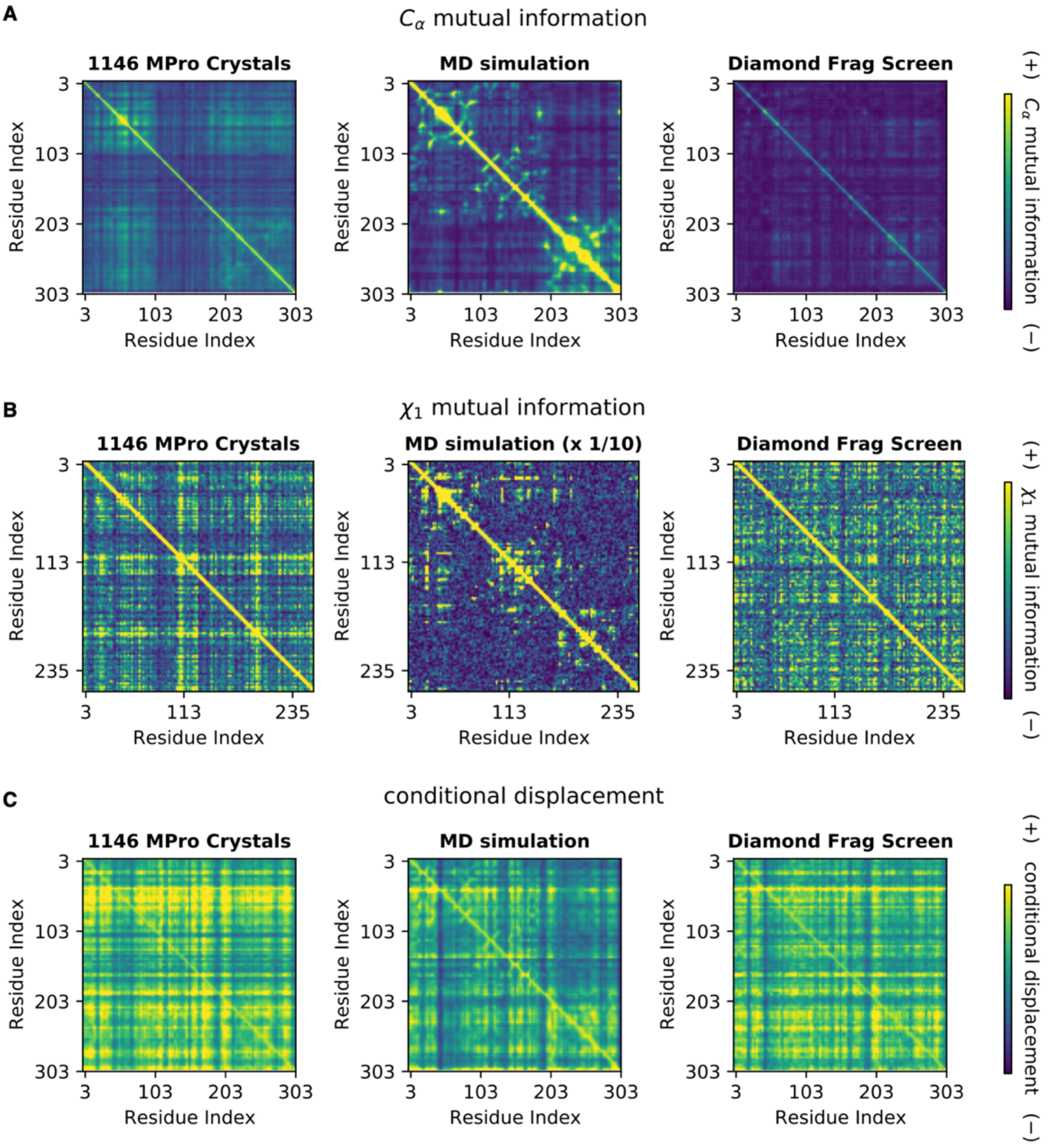


**Supplemental Figure 3. Alternative measures of correlated motion.** Reported are (**A**) the mutual information between $C_\alpha$ cartesian positions, (**B**) the mutual information between sidechain $\chi_1$ dihedral angles, and (**C**) the conditional displacement of $C_\alpha$ atoms (see *Methods and Materials* for definitions). Mutual information is able to capture non-Gaussian behavior, but requires more samples to converge than the covariance measure employed in the main text. The conditional displacement assumes Gaussian displacements, but can capture anisotropic motion, and therefore also requires additional data to converge. The MD simulation ensemble (middle column), consisting of 10,000 structures sampled across 100 μs of simulation time, shows the structure of the correlations present in the protein in the mutual information metrics (top two rows). In contrast, for the crystalline ensembles (rightmost and leftmost columns), the 1146 and 95 crystal structure, respectively, exhibit poorly converged digitized structure for all three measures reported in this figure. Note the MD simulation color scale zoomed out by a factor of 10 in the $\chi_1$ mutual information plot (central panel in B).

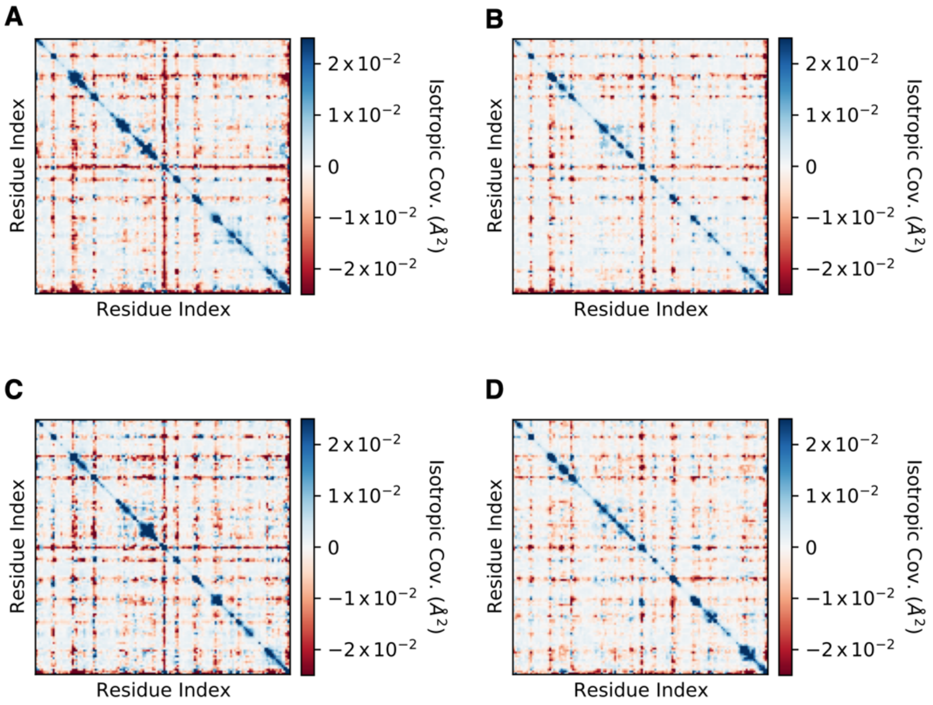


**Supplemental Figure 4. Ensembles fit to single crystal datasets do not reproduce the covariance structure provided by multiple crystalline datasets.** Because our automatic refinement procedure employed MD-driven simulated annealing and used standard crystallographic restraints that are effectively an MD forcefield, we tested the idea that the distribution of structures we observed in our crystallographic ensemble might simply be the result of the forcefield prior used during refinement, and not the experimental data themselves. We conducted ensemble refinement with *phenix* (*2*, *3*) on four datasets chosen to span the observed cell volumes, a structurally diverse and representative sample from our collection of $M^{pro}$ crystal data. The resulting ensembles are qualitatively different from the ensemble formed by the collection of all crystal structures. See Table S2 for cell volumes, ensemble size, and *R*-factors for these four datasets.

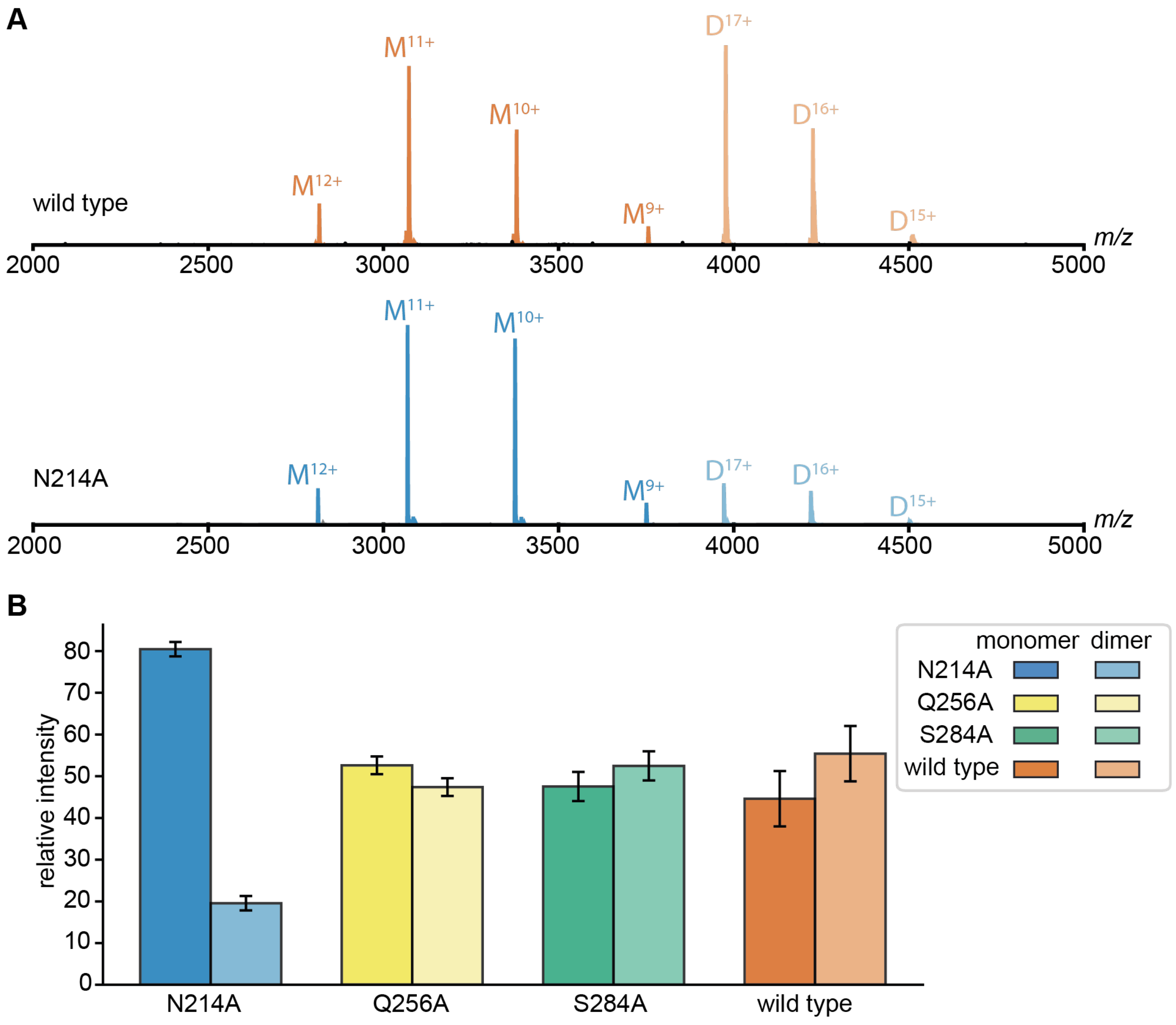


**Supplemental Figure 5. Native MS shows decreased dimer/monomer ratio for N214A variant as compared to wild type and other mutants.** (**A**) Representative native mass spectra of 2 µM wild type (top) and 2 µM N214A (bottom) showing distinct species distribution. N214A exhibits predominantly monomer peaks, whereas monomer and dimer are equally distributed in wild type. (**B**) Based on triplicate measurements, the bar chart shows the average relative intensities, proportional to the species concentration of monomers and dimers of all variants at 2 µM (see Supplemental Table 4). Error bars are standard errors.

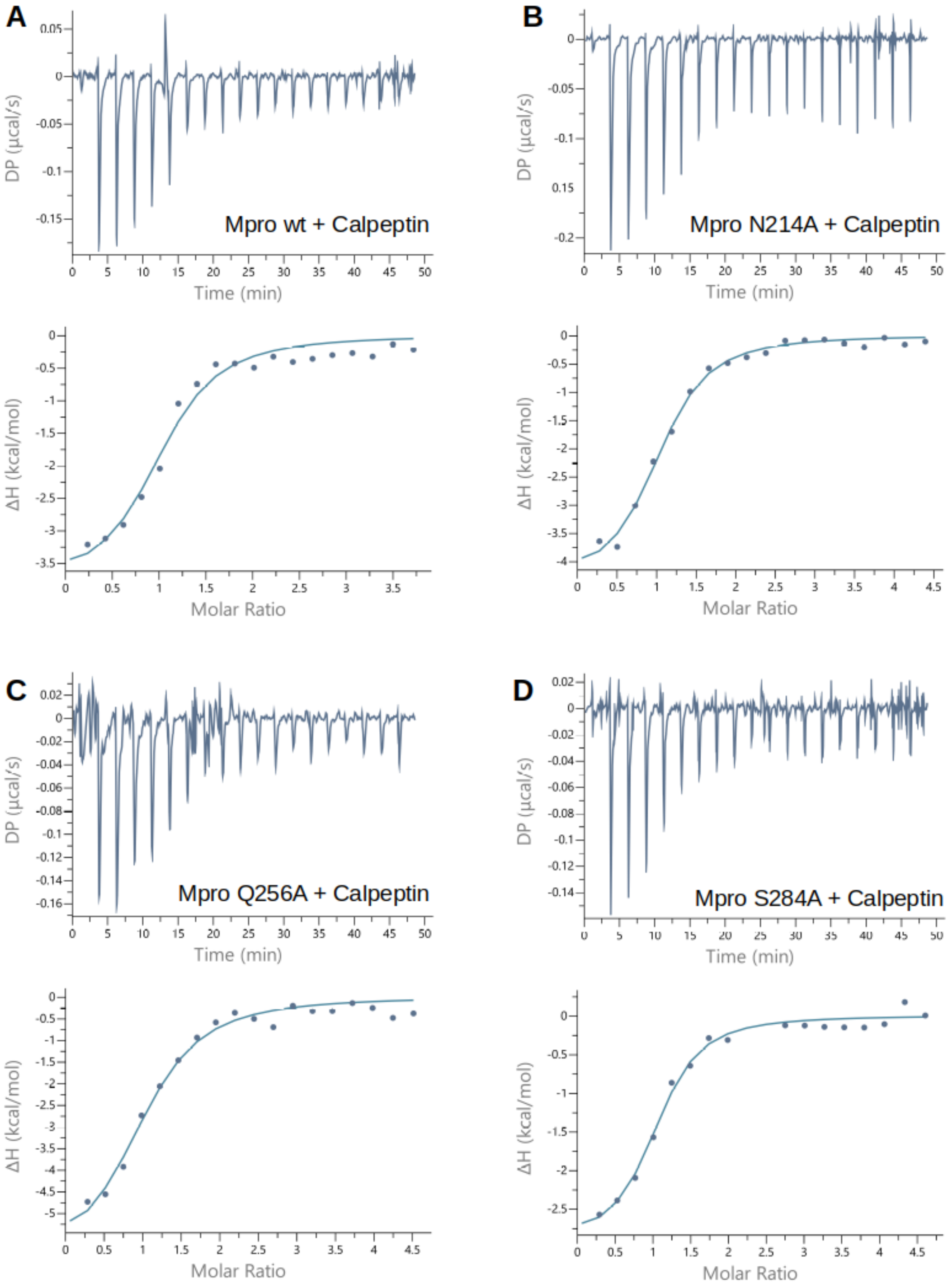


**Supplemental Figure 6. Isothermal titration calorimetry (ITC) shows ligand affinity is similar for all M$^{pro}$ variants studied.** Shown are the binding isotherm and fit for (**A**) wild type, (**B**) N214A, (**C**) Q256A and (**D**) S284A. Upper panels show the instrument power during the course of ligand injections. Lower panels show the evolved enthalpy per mol of injectant (calpeptin) against the ligand/protein molar ratio. Two points in the lower panel of (D) were rejected as outliers by the ITC software during the data analysis stage. The similarity of all binding isotherms and the ligand $K_D$ from Table 3 indicates a minor effect of the mutations studied on the ligand binding affinity. See Supplemental Table 3 for thermodynamic parameters obtain from isotherm fits.

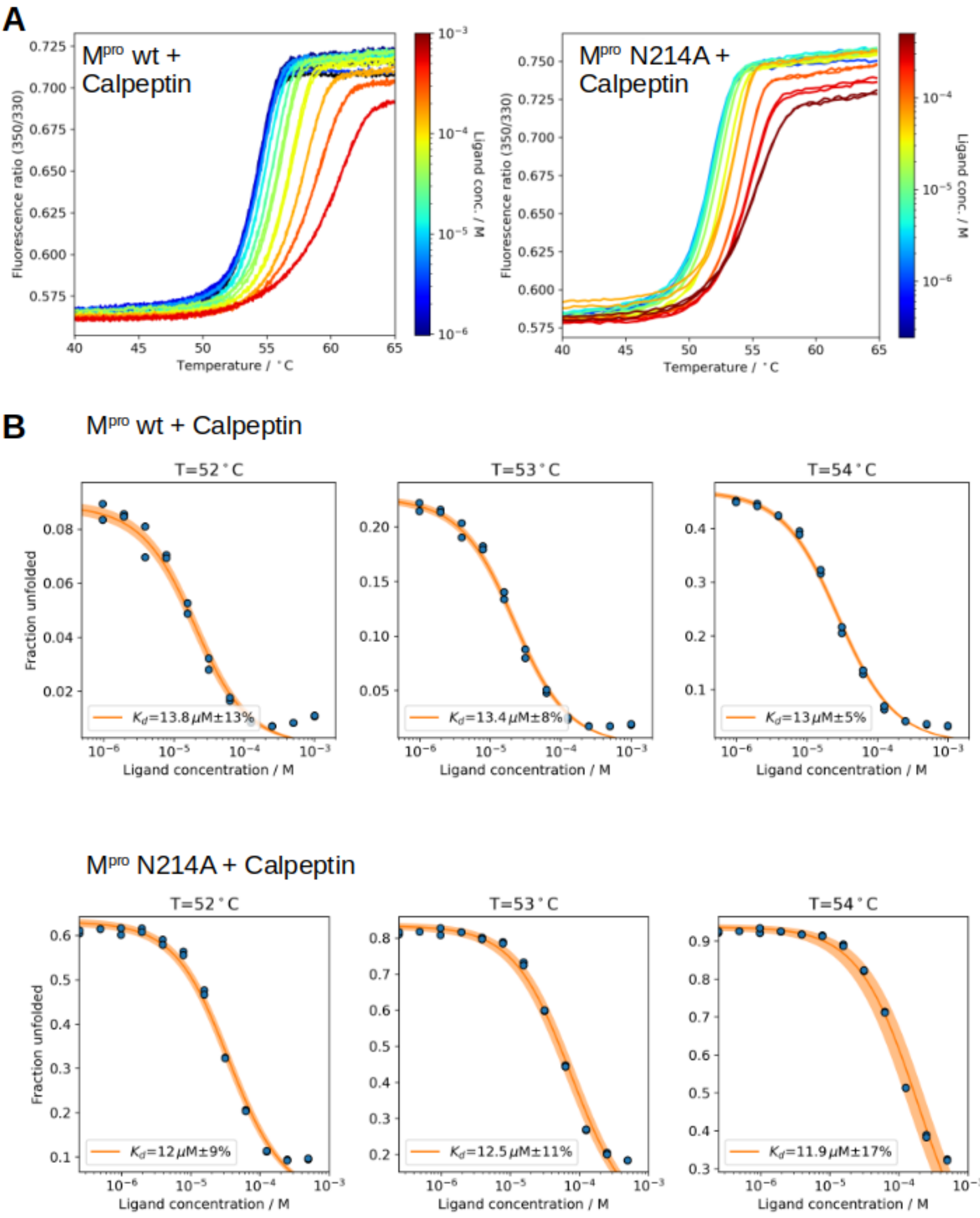


**Supplemental Figure 7. Nano differential scanning fluorimetry (nDSF) confirms ITC-derived binding affinity of M$^{pro}$ N214A.** To confirm the affinity of calpeptin for M$^{pro}$ N214A (ligand $K_D$), we conducted nDSF experiments. (**A**) Fluorescence ratio 350 nm/330 nm for nDSF binding studies of wild-type M$^{pro}$ and M$^{pro}$ N214A in the presence of increasing concentrations of calpeptin. **B**) Isothermal analysis for three selected temperatures, each showing the fraction of unfolded protein, determined by changes in intrinsic fluorescence (emission ratio at 350 nm/emission at 330 nm), as a function of ligand concentration. Fit dissociation constants (ligand $K_D$) determined from the curve fits are approximately 13 µM for the wild type and 10 µM for the N214A variant. The two tested proteins show slightly different melting temperatures of 54 °C (wild type) and 51 °C (N214A). Isothermal analysis can estimate the $K_D$ close to the melting temperature of the protein (*5*). Note the ligand affinity is reduced by about a factor of five in our nDSF measurement as compared $K_D$s determined from ITC (Supplemental Table 3); the higher temperature of the nDSF experiment is the most likely origin of this discrepancy, though we did not investigate this further. Nevertheless, the wild-type M$^{pro}$ and the N214A mutant ligand affinities are not distinguishable within the error of the nDSF measurement, demonstrating the N214A mutant's ligand binding capabilities are not greatly compromised. All plots shown here were generated with BioPhyPy (*6*).

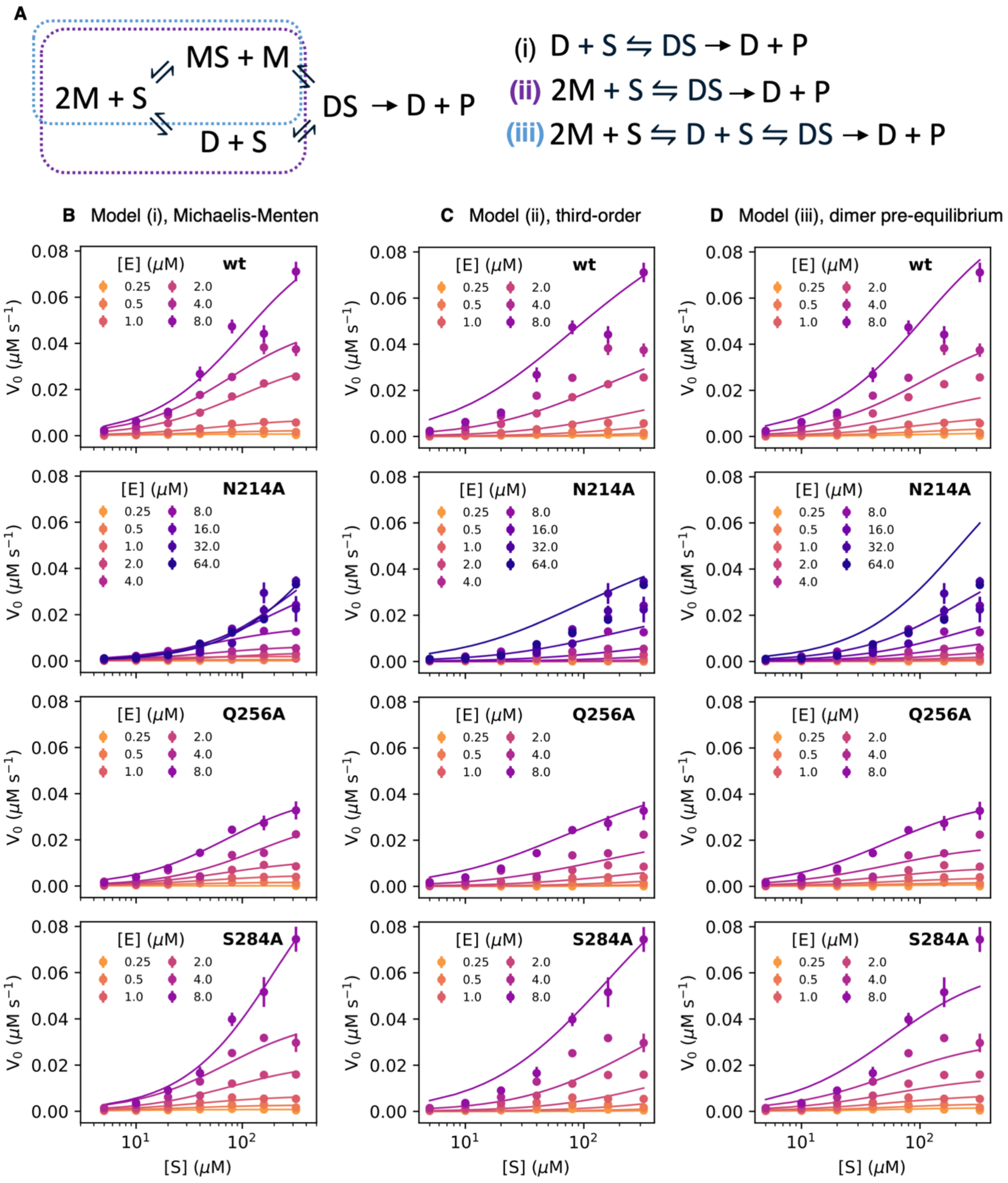


**Supplemental Figure 8. Enzyme kinetics models considered.** (**A**) Elaborated kinetic scheme for catalysis by M$^{pro}$ dimers. As this full scheme contains many fundamental rates that cannot be unambiguously determined, we assessed simplified models to explain observed initial reaction velocities $V_0$ as a function of total enzyme concentration [E] and substrate concentration [S]. Additional details and equations can be found in the *Materials and Methods.* (**B**) In model (i), each enzyme concentration is treated as an independent Michaelis-Menten scheme, as presented in the main text (Fig. 4). (**C**) In model (ii), we assume substrate equilibrates rapidly with both monomer and dimer species, which in turn equilibrate rapidly with one another (species inside purple box in (A)), leading to third-order association kinetics with a fit $K_{eq}$. This model provides a slightly worse fit, with a comparable interpretation from the fit parameters (Supplemental Table 5). Note that N214A is not well described by this model and the kinetic parameters are meaningless. (**D**) Finally, in model (iii), the monomer/dimer equilibration is assumed to be slow with respect to the rate of formation of a dimer-substrate complex, such that for each enzyme concentration a fixed concentration of dimers turns over substrate with Michaelis-Menten kinetics (species in blue box in (A)). This model fits the observed data reasonably well for all variants except N214A. While qualitatively different, the qualitative conclusions drawn from each model are consistent. See Supplemental Table 5 for fit parameters. Error bars are 95% confidence intervals.

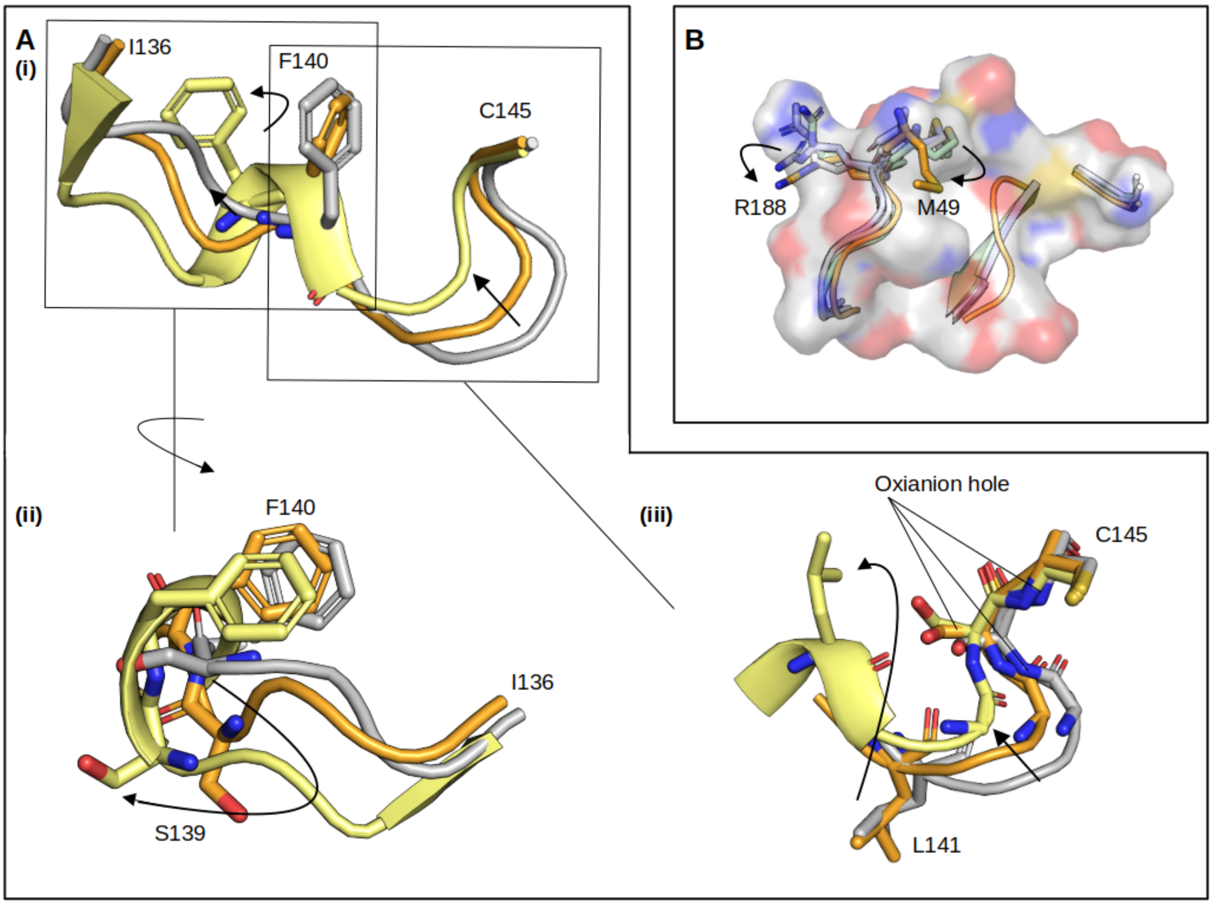


**Supplemental Figure 9. Comparison of the active site between M$^{pro}$ variants and the monomeric construct M$^{pro}$1-199.** Wild-type M$^{pro}$ (PDB: 7BB2) is depicted in grey, while the N214A variant (PDB: 9GI6) is orange and the monomeric M$^{pro}$1-199 construct (PDB: 7UJ9) in yellow (*7*, *8*). (**A**)(i) In the stabilized monomeric M$^{pro}$ structure, the oxyanion hole loop (I136 to C145) forms a $3_{10}$ helix (residues S139 to L141) not seen in the wild-type reference structure. While a clear $3_{10}$ helix was not found in the structure of the N214A mutant, we detected a loss in observable density and shift in mean position of residues L141 to G143, potentially reflecting an intermediate state between the monomer construct and wild type (see main text Fig. 5). (ii) Close-up view of the left side of panel (i), rotated for clarity. Amino acid S139 exhibits a side-chain flip coupled with realignment within the loop. (iii) Close-up view of the right side of the loop in panel (i), showing L141 occupying F140's original position to form a helix in the monomeric construct. (**B**) An alternative view of the M$^{pro}$ active site. Although the flexible residues R188 and M49 exhibit slight conformational shifts, the three different point mutations induce minimal changes to the active site architecture. Arrows highlight the changes that are observable, which are most dramatic in the N214A mutant.

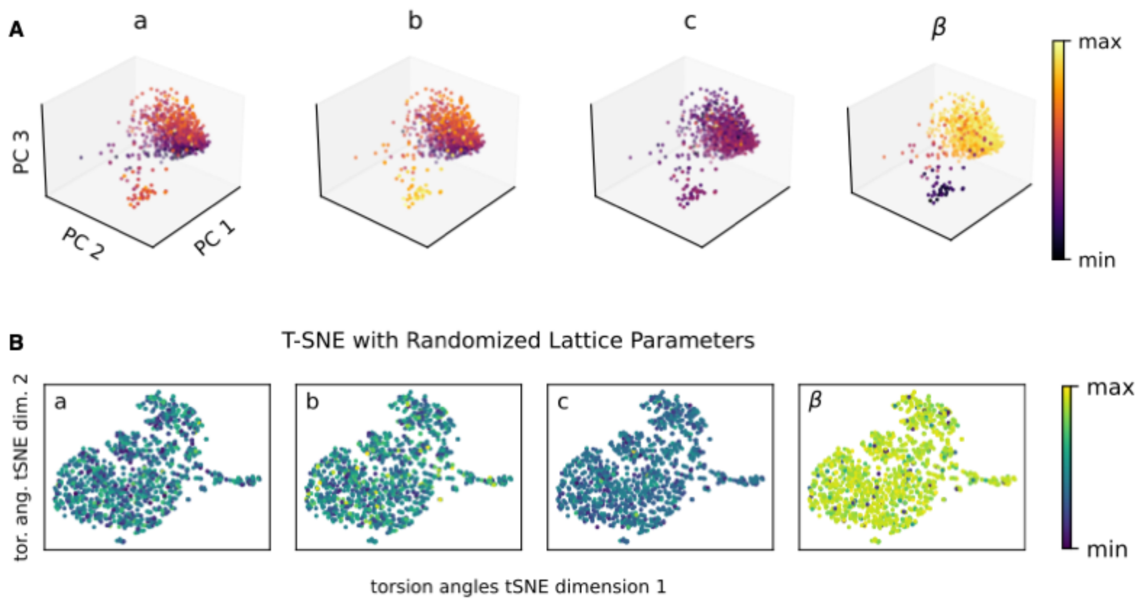


**Supplemental Figure 10. Controls for t-SNE analysis that support a link between protein structure and crystal lattice.** (**A**) PCA is presented as an alternative dimensionality reduction method to visualize the lattice-structure relationship. PCA was performed on the sine and cosine of each backbone dihedral angle, and three first (highest variance) principal components (PC1, PC2, PC3 respectively) were plotted. These components account for 19%, 8%, 6% of the total variance, respectively. The points are colored by the relative lattice parameter length, where the relevant lattice dimension is indicated above each plot. (**B**) The same t-SNE analysis as shown in main text Figure 6C, but with the lattice parameter values randomized. The lack of any structure is apparent.

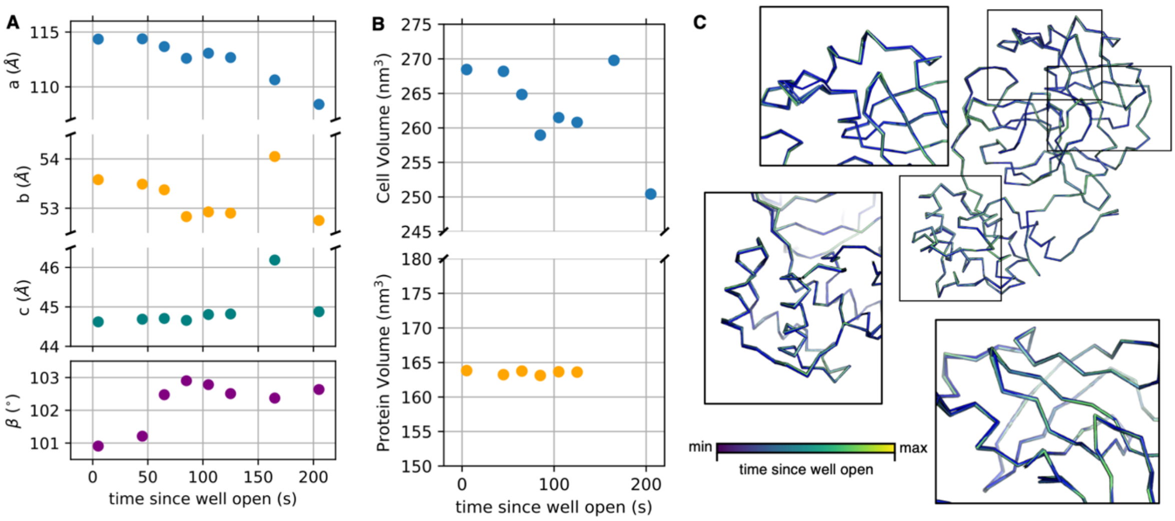


**Supplemental Figure 11. Delayed crystal fishing produces a continuous deformation of the M$^{pro}$ structure.** A single well of M$^{pro}$ crystals was opened and crystals were fished and frozen continuously. These crystals were then subjected to diffraction analysis and models were refined to the resulting data. (**A**) As a function of time since the well was opened, the cell parameters change continuously, (**B**) resulting in a contraction of the unit cell (top panel) but no appreciable change in the refined protein volume (bottom panel, protein volume determined by *3v* (*9*)). This implies that the change in cell volume is due to a loss of solvent as the crystals dry out. (**C**) The corresponding protein structures change continuously as a function of time since the well was opened. Shown are 6 structures from 5 seconds delay (blue) through 125 seconds delay (light green). Crystals fished at 165 and 205 seconds, the rightmost data points, did not produce sensible structures (*R*-factors >40%) and can be considered unreliable. The structural changes are similar to those seen when the ensemble of M$^{pro}$ crystals analyzed in the main text is sorted by unit cell volume (Figure 6B). We conclude that hydration changes, caused for example by delays during crystal fishing, account for a major factor in generating the diversity of structures we analyzed.
